# Intrinsic connectivity of the prefrontal cortex and striato-limbic system respectively differentiate Major Depressive from Generalized Anxiety Disorder

**DOI:** 10.1101/2020.05.19.105148

**Authors:** Xiaolei Xu, Jing Dai, Yuanshu Chen, Congcong Liu, Fei Xin, Xinqi Zhou, Feng Zhou, Emmanuel A Stamatakis, Shuxia Yao, Lizhu Luo, Yulan Huang, Jinyu Wang, Zhili Zou, Deniz Vatansever, Keith M Kendrick, Bo Zhou, Benjamin Becker

## Abstract

Major Depressive Disorder (MDD) and Generalized Anxiety Disorder (GAD) are highly prevalent and debilitating disorders. The high overlap on the symptomatic and neurobiological level led to ongoing debates about their diagnostic and neurobiological uniqueness. The present study aims to identify common and disorder-specific neuropathological mechanisms and treatment targets in MDD and GAD. The present study combined categorial and dimensional disorder models with a fully data-driven intrinsic network level analysis (Intrinsic Connectivity Contrast, ICC) to resting state fMRI data acquired in 108 partn = 35 and n = 38 unmedicated patients with first-episode GAD, MDD respectively and n=35 healthy controls). Convergent evidence from categorical and dimensional analyses revealed MDD-specific decreased whole-brain connectivity profiles of the medial prefrontal and dorsolateral prefrontal cortex while GAD was specifically characterized by decreased whole-brain connectivity profiles of the putamen and decreased communication of this region with the amygdala. Together, findings from the present data-driven analysis suggest that intrinsic communication of frontal regions engaged in executive functions and emotion regulation represent depression-specific neurofunctional markers and treatment targets whereas dysregulated intrinsic communication of the striato-amygdala system engaged in reinforcement-based and emotional learning processes represent GAD-specific markers and a promising treatment target.

## Introduction

Major depressive (MDD) and anxiety disorders are among the most prevalent and devastating disorders. Comorbidity represents the normative clinical course, with particular high rates between MDD and Generalized Anxiety Disorder (GAD) (lifetime comorbidity 70-90%) [1]. Together with the high comorbidity rates, a strong overlap in symptomatology, genetic and risk factors [1] led to a continuous debate about the nosological and neurobiological uniqueness of GAD and MDD [2].

Overarching symptom-based approaches suggest that both disorders share general affective distress accompanied by distinguishing features such as anhedonia (depression) and physiological hyperarousal (specific to anxiety disorders) [3]. Further evidence for common and distinct features is provided by case control studies that examined behavioral and neural dysregulations in MDD and GAD. Studies comparing either MDD or GAD patients with healthy controls revealed cognitive deficits in the domains of executive functions, complex attention, reward processing and social cognition [4] with initial evidence for separable alterations in emotion-specific processing bias and attributional style from studies directly comparing MDD and GAD patients [5].

In line with the behavioral dysregulations in domains strongly associated with the integrity of cortico-striato-limbic circuits qualitative and quantitative reviews covering functional magnetic resonance imaging (fMRI) studies suggest largely overlapping dysregulations in regional brain activation in this circuitry in both MDD and GAD patients. Relative to healthy controls some studies have reported increased responses in the amygdala, insula and dorsal anterior cingulate cortex, accompanied by decreased activity in the dorsal striatum, dorsolateral and medial prefrontal cortex (MPFC) in MDD patients during emotional and cognitive processing [6], however, a recent meta-analysis failed to determine robust task-related regional activation alterations [7]. Whereas several neuroimaging studies have examined anxiety patients and produced overwhelming, yet partly inconsistent results for dysfunctional processing in limbic and frontal regions [8], comparably few studies specifically focused on GAD. Examining emotional and cognitive processes these studies partly suggest exaggerated limbic reactivity in the context of both decreased as well as increased frontal activation [2].

Although meta-analyses of case control studies greatly advanced our knowledge of the pathological mechanisms that underlie MDD and GAD the high convergence of behavioral and neurobiological signatures has led to a continuing debate about the degree to which the clinical diagnosis of MDD and GAD reflect distinct neurobiological mechanisms [2]. Recent meta-analyses revealed only few differences in brain structural and functional indices between distinct diagnostic categories [9], suggesting that traditional case control designs are of limited ability to determine disorder-specific brain-based biomarkers and emphasizing the need for dimensional and transdiagnostic approaches while strictly controlling for confounding factors such as medication and disorder duration [10].

To date only a few neuroimaging studies rigorously employed this approach and included MDD and GAD patients across diagnostic categories. Task-based neuroimaging studies revealed common and distinct alterations between MDD and GAD patients in amygdala, cingulate cortex and dorsomedial prefrontal cortex activation, yet separable alterations in ventrolateral prefrontal cortex and anterior insula activation and amygdala functional connectivity with the specific regions identified depending on the task paradigm [11, 12]. Together with evidence for emotion-specific behavioral deficits this indicates that the observed alterations may vary as a function of emotional context and this resonates with current findings suggesting that task-independent assessment of intrinsic (resting state) brain function might represent a more general and robust neuroimaging-based biomarker [13].

To date two studies have combined a transdiagnostic and dimensional approach with resting state fMRI to determine common and specific neural signatures of MDD and GAD. Both studies focused on the intrinsic organization of a priori defined regions or networks of interest and demonstrated that subgenual anterior cingulate cortex and ventral striatum connectivity exhibited opposite associations with MDD versus GAD symptom load [14] and that resting-state connectivity between the limbic network and cortical regions specifically characterized patients with comorbid MDD and GAD [15]. However, the interpretation of these findings is limited by an a priori focus on brain systems previously identified in case control studies and hypothesis-free data-driven approaches may promote a more unbiased determination of common and disorder-specific alterations to inform neuropathological models as well as the development of disorder-specific treatment approaches. In fact the assessment of intrinsic brain functional connectivity has received increasing attention in psychiatric neuroimaging not only for the identification of neuropathological mechanisms but also the identification of therapeutic targets for focal brain stimulation [16]. Focal brain stimulation has been associated with changes that spread through the intrinsic networks of the brain and which outlast the actual stimulation period [17], including stimulation of the dorsolateral prefrontal cortex (DLPFC) which represents the primary and currently most effective target for non-invasive brain modulation in both MDD and GAD [18, 19]. In line with these findings the anti-depressive treatment efficacy of non-invasive transcranial DLPFC stimulation has been associated with effects in distal brain regions and re-organization of multiple intrinsic brain networks, suggesting an important role of network level effects for therapeutic efficacy [20]. The most efficient stimulation targets across invasive and non-invasive targets are linked by shared intrinsic networks, further emphasizing the therapeutic and mechanistic importance of network level effects [21]. However, across focal brain stimulation approaches treatment response critically relies on the specificity and connectivity of the stimulation site emphasizing the need for the identification of disorder-specific targets to promote therapeutic efficacy.

To identify common and disorder-specific neuropathological mechanisms and treatment targets for focal brain stimulation approaches the present study combined categorial and dimensional disorder models with a fully data-driven intrinsic network level analysis that operates independently of a priori assumptions (Intrinsic Connectivity Contrast, ICC) to resting state data acquired in 108 participants (n = 35 and n = 38 unmedicated patients with their first episode of GAD, MDD respectively, and n = 35 matched healthy controls).

## Methods and Materials

### Participants

108 participants were enrolled in this study including 35 unmedicated, first episode patients with generalized anxiety disorder (GAD) and 38 unmedicated patients with major depressive disorder (MDD) recruited at the Sichuan Provincial People’s Hospital and The Fourth People’s Hospital of Chengdu (Chengdu, China) and 35 healthy controls (HC) recruited by local advertisement. GAD and MDD diagnosis were determined by an experienced psychiatrist according to DSM-IV criteria (Sichuan Provincial People’s Hospital) or ICD-10 (Fourth People’s Hospital of Chengdu) and further confirmed by an experienced psychologist using Mini International Neuropsychiatric Interview (M.I.N.I.) for DSM-IV. All participants gave written informed consent after they were informed about the detailed study procedures and informed that they were allowed to withdraw from this study at any time without negative consequences. During the experimental assessments all participants underwent brain functional and structural MRI assessments (see e.g. also previous publication reporting findings from an empathy task-fMRI paradigm [11]), and were administered the Beck Depression Inventory II (BDI-II) and Penn State Worry Questionnaire (PSWQ) to determine MDD and GAD-symptom load, respectively. To ensure data validation and reduce the burden for participants, all of them were explicitly asked whether their current status (e.g. exhaustion, emotional state) allowed them to proceed with subsequent assessments (e.g. MRI assessments, questionnaires). The study and all procedures were fully approved by the local UESTC ethics committee and adhered to the latest revision of the Declaration of Helsinki.

All patients were unmedicated and had not previously received a diagnosis of or treatment for a psychiatric disorder. The diagnostic assessments were conducted during initial admission to the hospitals by experienced psychiatrists and suitable patients underwent the fMRI assessments during the period of further diagnostic clarification without receiving any treatment (<5 days after admission). The following exclusion criteria were applied to all participants, including controls: (1) history of or current episode of the following axis I disorders according to DSM criteria: post-traumatic stress disorder, PTSD, feeding and eating disorders, substance use disorders, bipolar disorder, and mania, (2) history of or current clinically relevant medical or neurological disorder, (3) acute (within six weeks before the assessments) or chronic use of medication, (4) acute suicidal tendencies, (5) contraindications for MRI assessments, (6) left handedness, and, (7) excessive motion during MRI assessment (head motion >3mm).

According to the exclusion criteria 10 subjects were excluded leading to a final sample of n=98 (HC = 33, GAD = 31, MDD = 34) for fMRI analyses. Specific reasons for exclusion are displayed in Supplementary Figure 1 and detailed diagnoses of GAD and MDD according to DSM criteria and M.I.N.I. reported in Supplementary information. To further account for subclinical co-morbid symptom load the categorical approach (comparing MDD, GAD and HC) was flanked by a dimensional analysis strategy examining associations with MDD and GAD symptom load in the entire sample (pooling the data from MDD, GAD, and HC). In each case the influence of the other symptom dimension was controlled using FSL PALM-alpha110 toolbox (https://fsl.fmrib.ox.ac.uk/fsl/fslwiki/PALM, Permutation Analysis of Linear Models, number of permutations = 10,000) including anxiety or depression symptom load as covariate in the analysis respectively. The variance inflation factor (VIF) was used to assess collinearity between anxiety and depression symptom load. The VIF = 2.4 indicated no problematic collinearity in this study [22, 23].

All healthy controls were without psychiatric disorders according to the M.I.N.I. interview. Several patients (and one HC) were too exhausted to continue with the questionnaires following the MRI assessments leading to the number of participants per group varying from 33 to 26 (BDI-II, PSWQ) and 32-23 (CTQ) respectively. Importantly, there was no significant difference in the number of participants remaining for analyses between groups (χ^2^ = 0.14, *p* = 0.93).

### Measurements

The Beck Depression Inventory II (BDI-II) and Penn State Worry Questionnaire (PSWQ) were employed to dimensionally assess the depressive and GAD symptom load in all participants. In addition, the Childhood Trauma Questionnaire (CTQ) was administered to control for potential effects of early life stress exposures on brain activation.

### MRI data acquisition

MRI data were collected using a 3.0 Tesla GE MR750 system (General Electric Medical System, Milwaukee, WI, USA). Scanning parameters are reported in Supplementary Information.

### MRI data preprocessing

Resting state fMRI (rsfMRI) data were preprocessed using FMRIB software library (FSL, http://www.fmrib.ox.ac.uk/fsl) routines combined with advanced independent component analysis (ICA-AROMA) [24]. The first five functional volumes were discarded to eliminate saturation effects and achieve steady-state magnetization. Preprocessing for the remaining volumes included nonbrain tissue removal using BET, slice timing, realignment, intensity normalization and smoothing with a Gaussian kernel of 6 mm full width at half maximum (FWHM) using FSL FEAT. Registration to high resolution structural images was carried out using FLIRT and further to standard space using FNIRT (nonlinear registration). Motion parameter, white matter and csf signal regression and band pass filter (0.01-0.1Hz) were performed using ICA-AROMA. Next, a strict noise reduction was conducted using the SPM12-based CONN fMRI connectivity toolbox to further remove noise from white matter signal, cerebrospinal fluid (CSF) signal, motion parameters and the first-order derivatives and apply linear detrending. The head movements for all participants were less than 3mm and no significant differences were found between groups in the mean frame-wise displacement (*F*_*2*,*95*_ = 0.99, *p* = 0.38).

### Intrinsic connectivity contrast and follow-up seed to voxel functional connectivity

To allow an unbiased determination of voxel-based global connectivity the intrinsic global connectivity contrast was computed. This novel measure utilizes a graph-theoretical approach similar to the degree index in complex networks but without the need for a priori assumptions on regions of interst or arbitrary thresholds by weighing voxel-wise connections with their average r^2^ [25]. The ICC index reflects the squared sum of mean connections of a given voxel with the rest of the brain, with higher values representing higher connectivity strength of a given voxel with every other voxel in the brain. To further determine the associated networks contributing to these differences, the regions revealing group differences in ICC analysis were used as seeds in further functional connectivity (FC) analysis [26]. Based on previous studies reporting altered amygdala connectivity in both GAD and MDD disorders [27, 28] and our recent finding that the insula exhibited altered functional connectivity with the amygdala in MDD rather than GAD patients, connectivity analyses focused on determining functional connectivity differences between regions identified in the ICC analysis and the probabilistically defined entire amygdala using a small volume correction (svc) [29].

Both ICC and seed to voxel functional connectivity analyses were implemented within the SPM12-based CONN toolbox. Results are reported with the threshold of whole brain cluster level *p*_FDR_ <.05 (*p*_uncorr_<.001) for the ICC and *p*_FWE_ <.05 (small volume correction) for FC.

## Results

### Demographic data and dimensional symptom load

Participants in MDD, GAD and HC groups were well matched in age (*p* = 0.17), gender distribution (*p*(Χ^2^) = 0.11) and education level (*p* = 0.54). Symptom load and early life stress were analyzed using Univariate ANOVA. Results revealed significant main effects of group for depressive symptom load (BDI-II, *F*_*2*,*89*_ = 83.93, *p*<0.001, η^2^_p_ = 0.65), GAD symptom load (BDI-II, *F*_*2*,*89*_ = 83.93, *p*<0.001, η^2^_p_ = 0.65) and early life stress (CTQ, *F*_*2*,*83*_ = 8.74, *p*<0.001, η^2^_p_ = 0.17). Post-hoc analyses indicated that depressive symptom load was higher in both GAD and MDD patients compared to HC, and in MDD compared to GAD patients. GAD symptom load and early life stress were significantly higher in both patient groups relative to HC, but not significantly different between the two patient groups (see table 1 for details).

**Table 1.** Demographics, symptom load and early life stress.

| Group | HC | GAD | MDD |  |  |  |  |
| --- | --- | --- | --- | --- | --- | --- | --- |
|  | N = 33 | N = 31 | N = 34 |  |  |  |  |
| Male | N = 12 | N = 15 | N = 8 |  |  |  |  |
|  | (36%) | (48%) | (24%) |  |  |  |  |
|  | Mean(SEM) | Mean(SEM) | Mean(SEM) | *F* | GAD vs HC | MDD vs HC | GAD vs MDD |
| Age(years) | 26.79(1.46) | 30.74(1.51) | 28.18(1.44) | $F_{2,95}=1.82$ | >0.18 | >0.18 | >0.18 |
| Education | 14.15(0.62) | 14.32(0.68) | 13.38(0.63) | $F_{2,90}=0.62$ | >0.92 | >0.92 | >0.92 |
| (years) | (N = 33) | (N = 28) | (N = 32) |  |  |  |  |
| PSWQ | 39.49(1.63) | 58.08(1.84) | 63.00(1.63) | $F_{2,89}=56.94$ | <0.001 | <0.001 | =0.144 |
|  | (N = 33) | (N = 26) | (N = 33) |  | ** | ** |  |
| BDI-II | 5.21(1.49) | 23.15(1.68) | 32.18(1.49) | $F_{2,89}=83.93$ | <0.001 | <0.001 | <0.001 |
|  | (N = 33) | (N = 26) | (N = 33) |  | ** | ** | ** |
| CTQ | 43.00(1.98) | 50.30(2.34) | 54.71(2.01) | $F_{2,83}=8.74$ | =0.058 | <0.001 | =0.472 |
|  | (N = 32) | (N = 23) | (N = 31) |  |  | ** | ** |

PSWQ = Penn State Worry Questionnaire; BDI-II = Beck depression Inventory II; CTQ = Childhood Trauma Questionnaire; Given that some participants did not completed all questionnaires (details see also: Exclusion criteria, initial quality assessments and final sample) the number of subjects that indicated the respective analysis is reported for each measure. \*\* $p<.005$

### Intrinsic connectivity contrast

The ICC functional connectivity maps were used to examine group differences in CONN (see Supplementary Figure 2 for ICC maps in each group). The categorical analysis employed a one-way ANOVA with group as between-subject factor and revealed significant group differences in right medial prefrontal cortex (R_MPFC, x/y/z: 16/46/44, *k* = 34, *p*_FDR_ = 0.035, **Fig. 1A**), right putamen (x/y/z: 22/0/-4, *k* = 28, *p*_FDR_ = 0.041, **Fig. 1B**), and left dorsolateral prefrontal cortex (L_DLPFC, x/y/z: -40/24/28, *k* = 41, *p*_FDR_ = 0.027, **Fig. 1C**). To determine specific alterations between groups parameter estimates from the three clusters were extracted for post-hoc analyses. Results indicated that specifically R_MPFC connectivity was decreased in MDD but not GAD patients compared to HC (**Fig. 1D**) and the right putamen connectivity was decreased in GAD patients compared to both MDD and HC groups (**Fig. 1E**). In addition, both GAD and MDD patients exhibited decreased L_DLPFC connectivity compared to HC (**Fig. 1F**).

**Fig. 1.**
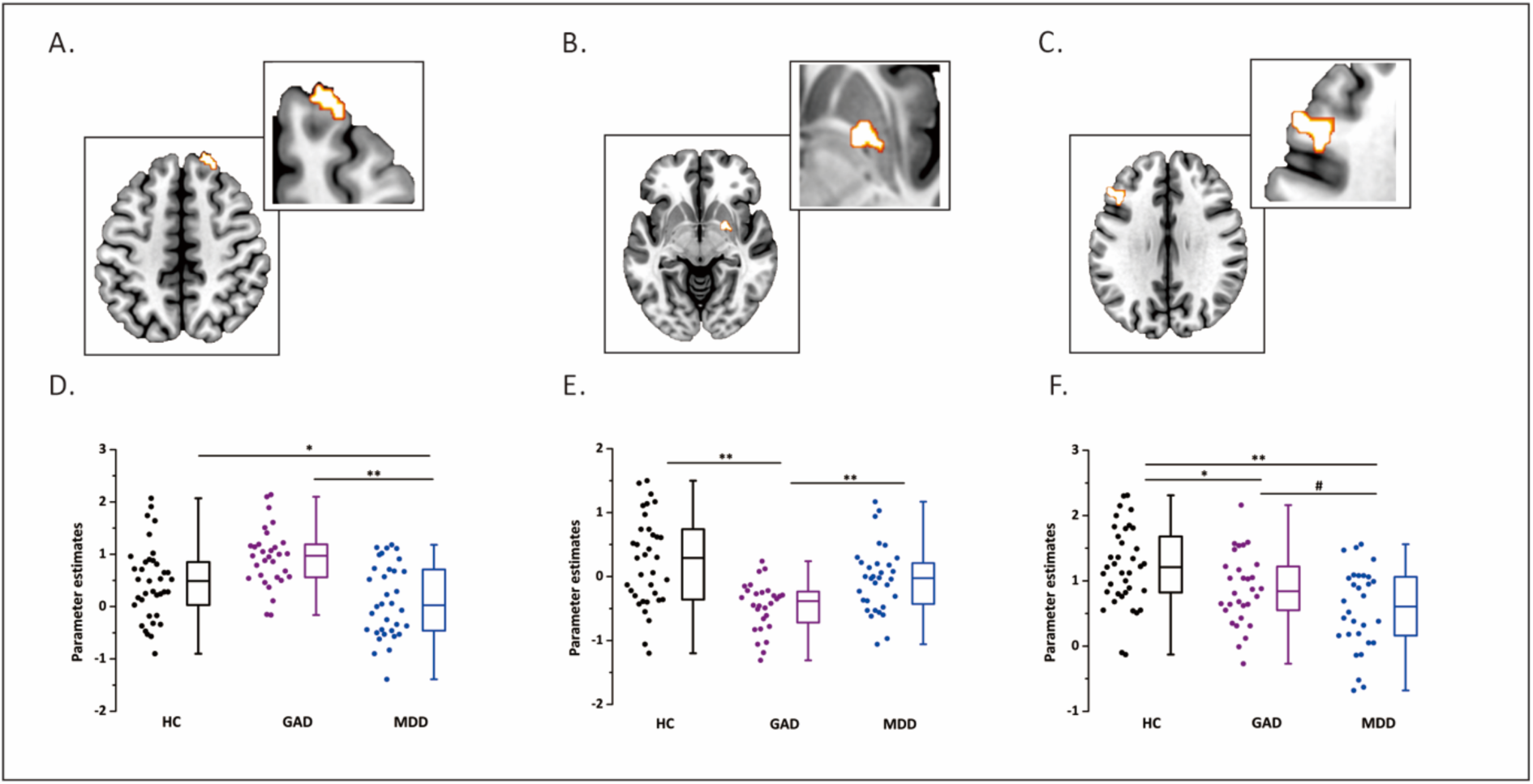
Brain areas exhibiting alterations in ICC analysis. A. right medial prefrontal cortex (R_MPFC); B. right putamen; C. left dorsomedial prefrontal cortex (L_DLPFC); D. group differences in R_MPFC; E. group differences in right putamen; F. group differences in L_DLPFC. R_MPFC = right medial prefrontal cortex, L_DLPFC = left dorsolateral prefrontal cortex. For visualization, statistical maps are displayed with a threshold of p<0.005 uncorrected.

### Follow-up seed-to-voxel functional connectivity

To further determine common and disorder-specific alterations in amygdala connectivity a functional connectivity analysis was performed using seeds from the regions which were found in ICC (R_MPFC, right putamen, L_DLPFC). One-way ANOVA with group as between subject factor found a main effect of group in right amygdala (centromedial, x/y/z: 29/-12/-9, *p*_svc-FWE_ < 0.05, **Fig. 2A**) when using the right putamen as seed. Post-hoc analysis indicated that both GAD and MDD patients exhibited decreased connectivity between putamen and amygdala compared to HC (**Fig. 2B**).

**Fig. 2.**
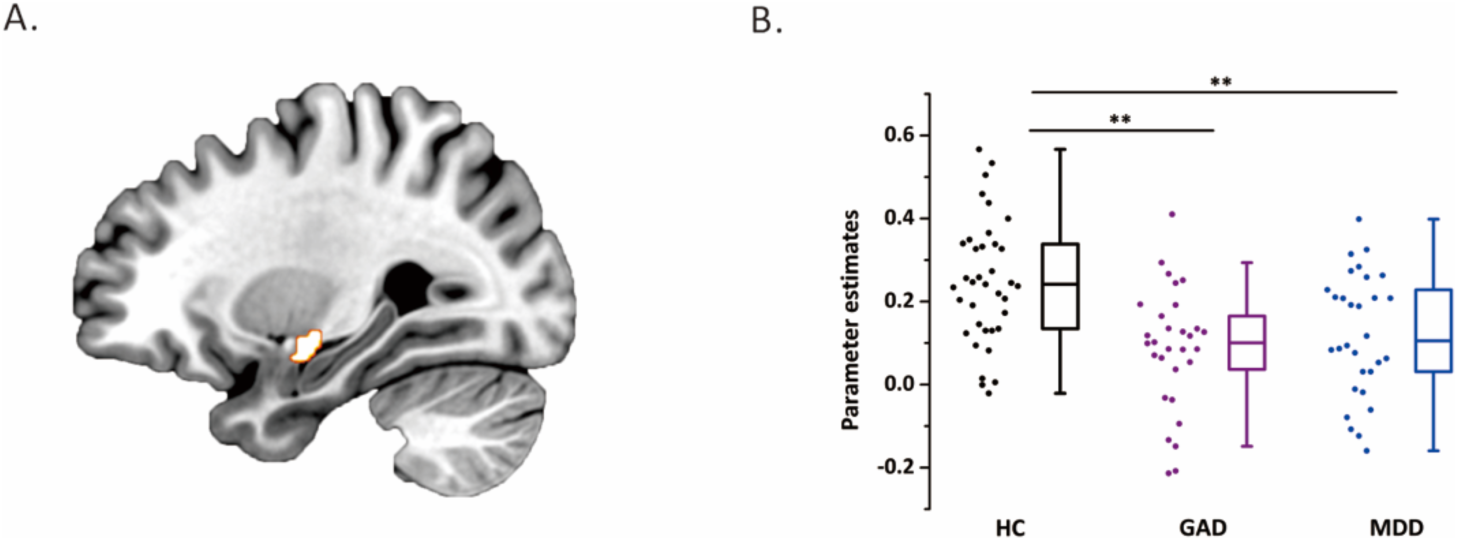
Brain regions showed aberrant functional connectivity with seeds from ICC. A. Altered right putamen – right amygdala (centromedial) connectivity and B. post – hoc group differences. ICC = intrinsic connectivity contrast.

### Dimensional analyses: associations between intrinsic network level indices and symptom load

Results from the dimensional analyses confirmed that depressive symptom load was negatively associated with ICC of the R_MPFC (r = −0.184, *p* = 0.038, **Fig. 3A**) and L_DLPFC (r = −0.376, *p* = 0.0002, **Fig. 3B**) while controlling for GAD symptom load, and that GAD symptom load was negatively associated with ICC of the right putamen (r = −0.190, *p* = 0.034, **Fig. 3C**) after controlling for depressive symptom load. On the functional connectivity level, connectivity strengths between putamen and amygdala were negatively correlated with GAD symptom load (r = −0.216, *p* = 0.015, **Fig. 3D**) after controlling for depressive symptom load.

**Fig. 3.**
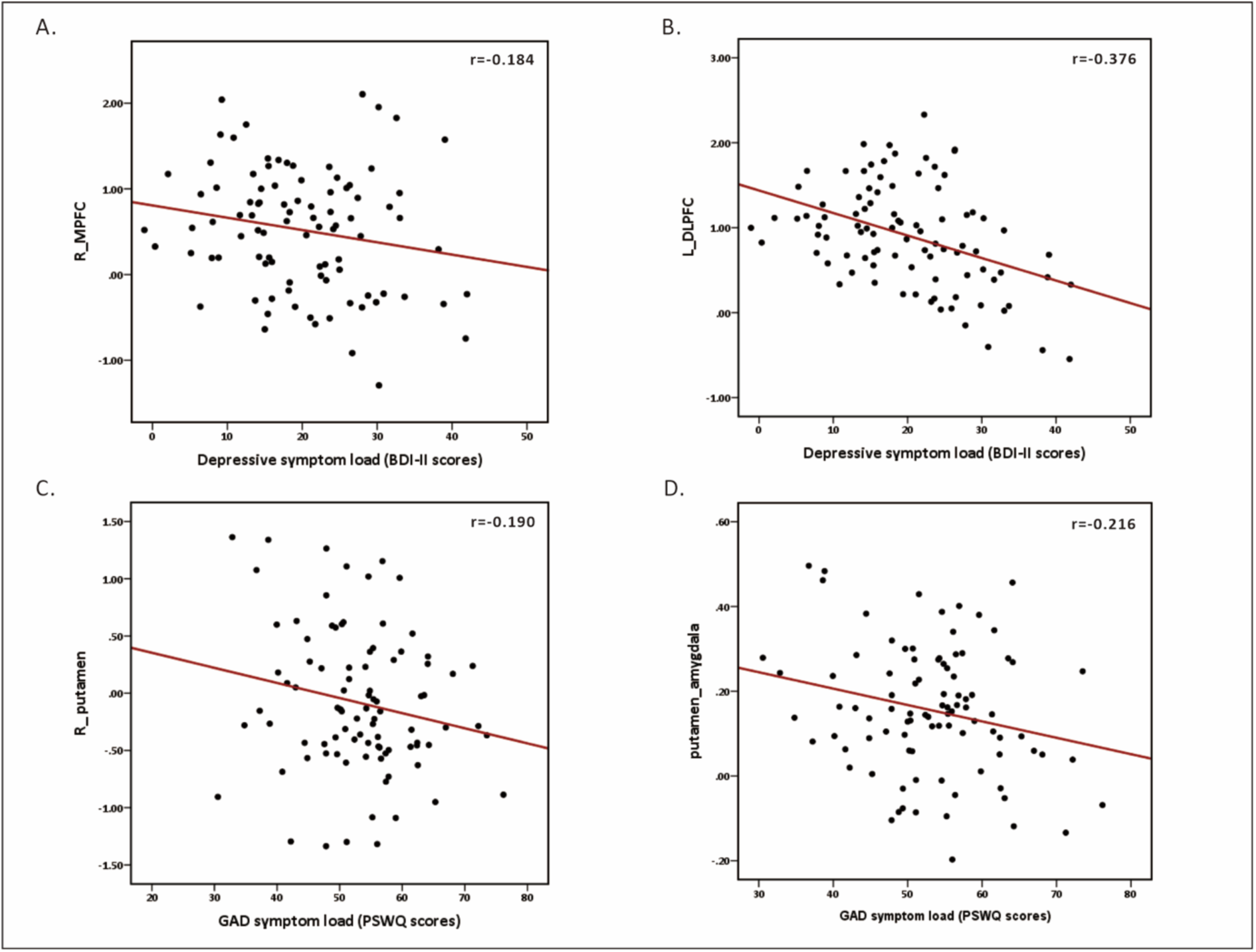
Associations between A. the right MPFC and depressive symptom-load; B. the left DLPFC and depressive symptom load; C. the right putamen and GAD symptom load; and D. the right putamen-right amygdala connectivity and GAD symptom load. Vertical axis reflects parameter estimates of corresponding brain areas. \**p*<.05, \*\**p*<.005. MPFC = medial prefrontal cortex, DLPFC = dorsolateral prefrontal cortex.

### Functional characterization of the altered brain areas

NeuroSynth decoding revealed that the highest correlated terms for R_MPFC were predominantly referring to cognitive processing and default mode network (**Fig. 4A**). For the right putamen, the highest correlated terms were gains, losses, learning and reward (**Fig. 4B**). For major depression, executive control, and working memory were highly correlated with the L_DLPFC (**Fig. 4C**).

**Fig. 4.**
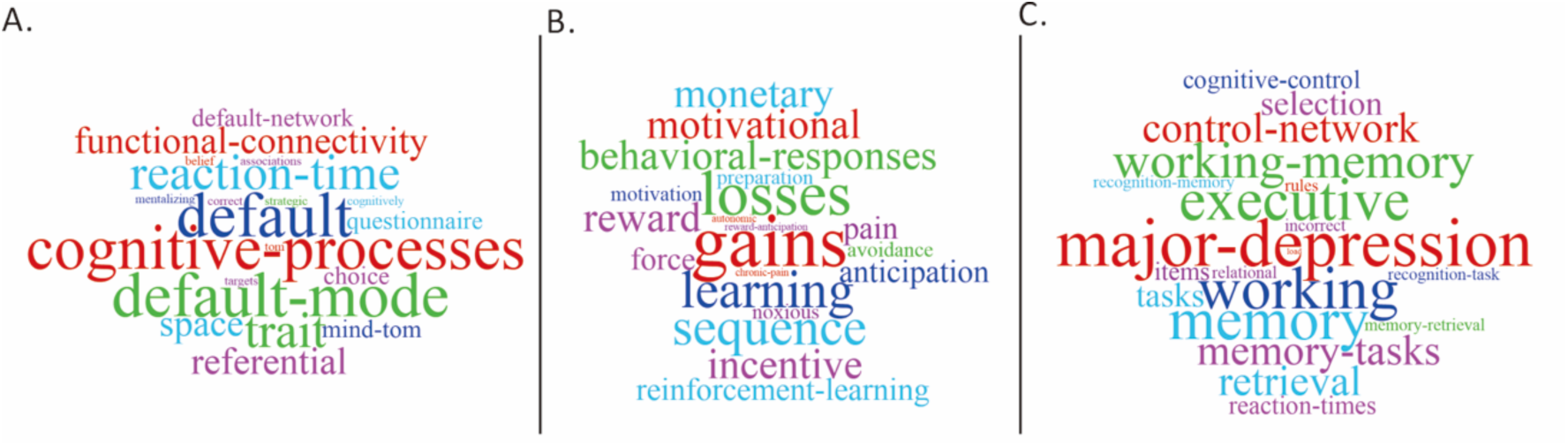
NeuroSynth decoding of A. right MPFC, B. right putamen, and C. left DLPFC. MPFC = medial prefrontal cortex, DLPFC = dorsolateral prefrontal cortex.

## Discussion

The present study applied for the first time a hypothesis-free fully data-driven approach in combination with a categorical and dimensional disorder model to determine common and disorder-specific alterations in whole brain intrinsic network connectivity in unmedicated, first-episode MDD and GAD patients. The categorical analysis approach demonstrated that MDD patients specifically exhibited decreased whole brain connectivity of the right MPFC compared to both HC and GAD, while GAD patients exhibited decreased right putamen whole brain connectivity relative to both other groups suggesting disorder specific neurofunctional deficits. In contrast, both patient groups demonstrated decreased DLPFC whole brain connectivity relative to HC, with pronounced deficits in MDD relative to GAD, and reduced putamen-amygdala connectivity, indicating common neurofunctional deficits in MDD and GAD. Dimensional analyses further confirmed symptom-specific alterations such that the strengths of whole brain connectivity alterations in the MPFC and DLPFC were associated with depressive symptom load whereas whole brain putamen and putamen-amygdala communication were associated with GAD symptom load. Together these findings provide evidence for a separable neurofunctional basis of the disorders, with MDD being characterized by deficient whole-brain connectivity of frontal regions and GAD being characterized by deficits in dorsal striatum whole-brain connectivity and functional communication of this region with the amygdala.

Intrinsic connectivity deficits in frontal regions including the MPFC and DLPFC have been repeatedly reported in depressive disorders and previous studies employing similar data-driven approaches demonstrated reduced global brain connectivity within the MPFC, ventromedial prefrontal cortex (VMPFC), anterior cingulate cortex (ACC) and DLPFC in MDD patients [30] relative to healthy control groups. In line with the functional characterization of the identified region, previous studies have demonstrated that the MPFC plays an important role in cognitive processes, including social cognition and emotion regulation [31], and represents a core hub of the brain’s anterior default mode system involved in self-referential processing, autographic memory and social cognition [32]. Structural and task-based functional MRI studies have consistently shown reduced brain volume and attenuated engagement of the MPFC during cognitive processes and emotion regulation in depression [30, 33] and a prospective study in patients with remitted depression suggests that attenuated MPFC reactivity to mood provocation may represent a risk factor for relapse [34]. Although previous meta-analytic approaches reported decreased emotion regulation associated MPFC engagement in anxiety disorders [35], studies focussed specifically on GAD have revealed somewhat inconclusive results [2].

Likewise, numerous studies have reported deficient DLPFC activation and connectivity during cognitive and emotional processing in both depressive and anxiety disorders [36], with less consistent results in GAD [2]. Although the results from the categorical analysis revealed that both MDD and GAD exhibit reduced left DLPFC whole brain connectivity relative to controls, deficits were more pronounced in MDD patients and specifically associated with depressive symptom load in the entire sample. In line with the functional characterization of this region, the DLPFC is a key region for executive functions and explicit emotion regulation [37] which have been consistently found to be impaired in MDD. In addition, non-invasive stimulation of the left DLPFC has been successfully applied in the treatment of MDD [38].

In contrast to depression-specific alterations in frontal regions, whole-brain connectivity of the putamen and its with the amygdala were found to be specifically associated with GAD symptom load. Both the functional characterization of this region and previous studies indicate an important role of the putamen in reinforcement (reward/punishment) learning and alterations in its responses have been found in anxiety populations during processing of reward-related information including monetary gains or losses [39, 40]. GAD patients have repeatedly been shown to exhibit deficient reinforcement-based learning with the degree of deficit observed being associated with both anxiety symptoms as well as punishment-related putamen activation [39, 41]. The amygdala conveys important information on both negative and postive signals in the environment, and critically contributes to the formation of emotional memories. The weakened connectivity between amygdala and putamen may therefore reflect an impaired integration of emotional experience with memory, or an inefficiency in utilizing recall of emotional memories as a tool to regulate or cope with emotional experience [27] promoting excessive and uncontrollable worry leading to impaired goal directed decision-making in GAD [42].

With respect to the diagnostics it is noteworthy that, although the categorical diagnostics by experienced clinical doctors and the structured M.I.N.I. interview clearly differentiated between the MDD and GAD patients the groups did not differ in the self-reported GAD symptom levels as assessed by the PSWQ suggesting a limited sensitivity of the self-reported (dimensional) measure to differentiate between these diagnostic categories.

Together, findings from the present data-driven study suggest that deficient intrinsic communication of frontal regions, specifically the MPFC and DLPFC which are strongly engaged in executive functions and emotion regulation, represent disorder-specific neurofunctional markers and treatment targets for MDD. On the other hand, impaired intrinsic communication of subcortical regions, specifically the putamen and its connections with the amygdala which are strongly engaged in reinforcement-based learning and emotional memory integration, characterize GAD and might represent promising targets for the treatment of this disorder.

## Authors contribution

XX, BB, and KMK designed the experiment. XX prepared the study protocols and procedures. BZ, JD, ZZ, YH, and JW performed the clinical assessments. XX, YC, CL, and FX acquired data. XX, FZ, XZ, EAS, LL, BB, and SY analyzed the data. XX, BB, KMK, and EAS interpreted the data and drafted the manuscript. All authors commented on and gave final approval to the final version of the manuscript.

## Acknowledgements

This work was supported by the National Key Research and Development Program of China (Grant No. 2018YFA0701400), National Natural Science Foundation of China (NSFC, No 91632117, 31530032, 31800961); Fundamental Research Funds for Central Universities (ZYGX2015Z002), Science, Innovation and Technology Department of the Sichuan Province (2018JY0001), Sichuan Science and Technology Program (2018JY0361), China Postdoctoral Science Foundation (2018M633336), Science and Technology Department of Sichuan Province, China (2017JY0031).

