## Supplementary Materials for "Intrinsic connectivity of the prefrontal cortex and striato-limbic system respectively differentiate Major Depressive from Generalized Anxiety Disorder"

**Exclusion criteria, initial quality assessments and final sample**

Study exclusion criteria, inspection of data quality and the independent diagnostic interview led to exclusion of n = 10 subjects due to technical issues during fMRI data collection (n=1), MDD and GAD diagnosis not validated by the M.I.N.I. or presence of a co-morbid previous or current disorder in the M.I.N.I. according to the exclusion criteria: PTSD (n=2), OCD (n = 1) (as primary diagnosis, GAD not confirmed by the M.I.N.I.), substance use disorder (n = 1), bulimia nervosa (n = 1), agoraphobia (n = 1), (as primary diagnosis, MDD diagnosis not confirmed in the M.I.N.I.), acute suicidal tendencies (n = 1), and mania (n = 2). The final sample for all fMRI analyses included HC = 33, GAD = 31, and MDD = 34 (total n = 98). All patients in the GAD group received the primary diagnosis GAD, all patients in the MDD group received the primary diagnosis MDD according to DSM criteria across the diagnostic assessment. Given the high prevalence of unipolar depressive and anxiety-associated disorders in GAD or MDD, respectively, a secondary diagnosis in these disorders that was additionally determined by the M.I.N.I. interview was thus not considered as exclusion criterion. N = 16 patients in the GAD (total n = 31) group and n = 17 (total n = 34) patients in the MDD group did not exhibit an additional psychiatric diagnosis. The following secondary diagnoses were determined by the M.I.N.I. interview: social phobia (GAD, n = 3, MDD, n = 3), obsessive compulsive disorder (GAD, n = 2; MDD, n = 2), panic disorder (GAD, n = 5; MDD, n = 1), agoraphobia (GAD, n = 5; MDD, n = 4), MDD (n = 7 in the GAD group), GAD (n = 6 in the MDD group).

### MRI data acquisition

MRI data were collected using a 3.0 Tesla GE MR750 system (General Electric Medical System, Milwaukee, WI, USA). Functional images were acquired within a single run using T2*-weighted echo planar imaging (sequence parameters, TR = 2000ms, TE = 30ms, flip angle = 90°, acquisition matrix = 64 × 64, thickness = 3.4mm, FOV = 240 × 240 mm, gap = 0.6mm, slices = 39). High-resolution whole-brain T1-weighted images were additionally obtained to exclude subjects with apparent brain pathologies and to improve normalization of the functional time series (sequence parameters, TR = 6ms, TE = minimum, flip angle = 9°, acquisition matrix = 256 × 256, thickness = 1mm, FOV = 256 × 256mm, slices = 156).

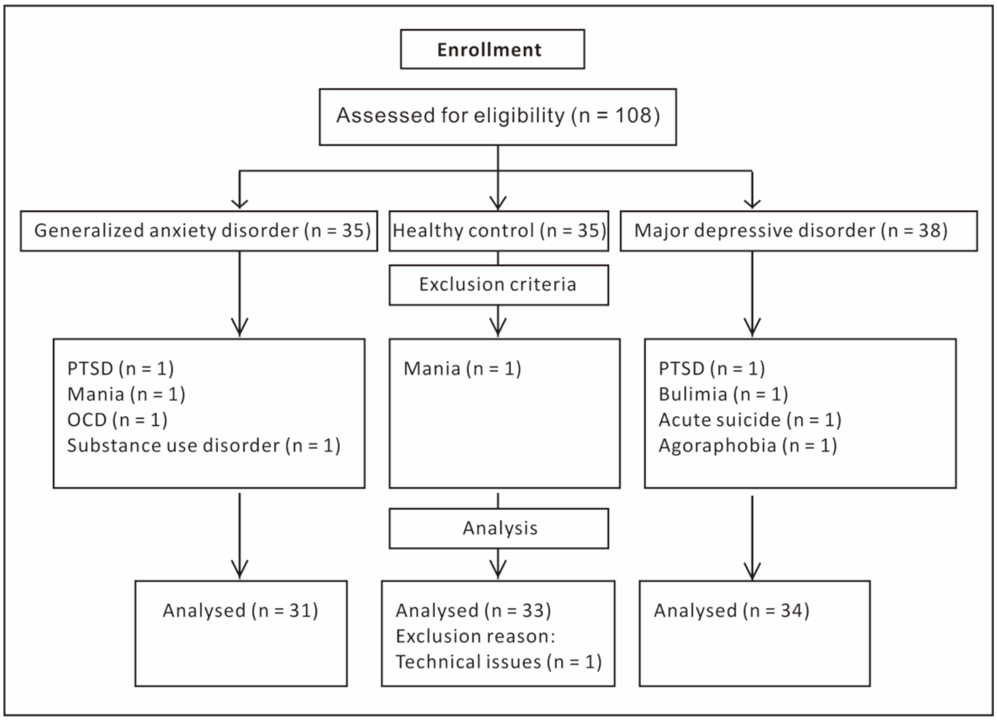


**Figure 1** CONSORT FLOW diagram

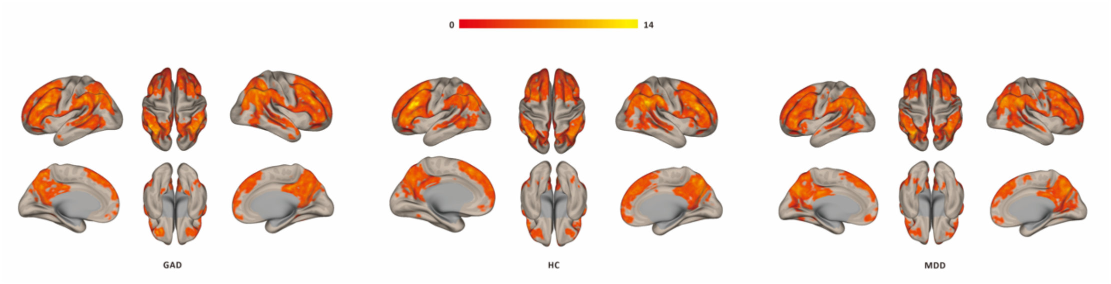


**Figure 2** Within-group intrinsic connectivity contrasts (ICC) maps.
